# Metformin Suppresses Monocyte Immunometabolic Activation by SARS-CoV-2 and Spike Protein Subunit 1

**DOI:** 10.1101/2021.05.27.445991

**Authors:** Theodore J. Cory, Russell S. Emmons, Johnathan R. Yarbro, Kierstin L. Davis, Brandt D. Pence

**Affiliations:** Department of Clinical Pharmacy and Translational Science, College of Pharmacy, University of Tennessee Health Science Center, Memphis, TN 38163; College of Health Sciences, University of Memphis, Memphis, TN 38152; Department of Medicine, University of Tennessee Health Science Center, Memphis, TN 38163; Center for Nutraceutical and Dietary Supplement Research, University of Memphis, Memphis, TN 38152

## Abstract

A hallmark of COVID-19 is a hyperinflammatory state that is associated with severity. Various anti-inflammatory therapeutics have shown mixed efficacy in treating COVID-19, and the mechanisms by which hyperinflammation occurs are not well understood. Previous research indicated that monocytes, a key innate immune cell, undergo metabolic reprogramming and produce inflammatory cytokines when stimulated with SARS-CoV-2. We hypothesized that binding by the viral spike protein mediates this effect, and that drugs which regulate immunometabolism could inhibit the inflammatory response in monocytes. Monocytes stimulated with recombinant SARS-CoV-2 spike protein subunit 1 showed a dose-dependent increase in glycolytic metabolism that was associated with production of pro-inflammatory cytokines including interleukin-6 and tumor necrosis factor-α. This response was dependent on hypoxia-inducible factor-1α, as chetomin inhibited glycolysis and cytokine production. Inhibition of glycolytic metabolism by 2-deoxyglucose (2-DG) or glucose deprivation also inhibited the glycolytic response, and 2-DG strongly suppressed cytokine production. Glucose-deprived monocytes rescued cytokine production by upregulating oxidative phosphorylation, an effect which was not present in 2-DG-treated monocytes due to the known effect of 2-DG on suppressing mitochondrial metabolism. Finally, pre-treatment of monocytes with metformin strongly suppressed spike protein-mediated cytokine production in monocytes, and abrogated glycolytic and mitochondrial metabolism. Likewise, metformin pre-treatment blocked cytokine induction by SARS-CoV-2 strain WA1/2020 in direct infection experiments in monocytes. In summary, the SARS-CoV-2 spike protein induces a pro-inflammatory immunometabolic response in monocytes that can be suppressed by metformin, and metformin likewise suppresses inflammatory responses to live SARS-CoV-2. This has potential implications for the treatment of hyperinflammation during COVID-19.

## Introduction

The ongoing coronavirus disease 2019 (COVID-19) pandemic has presently more than 3 million lives worldwide as of mid-April 2021 ^1^. COVID-19 is caused by a novel highly pathogenic coronavirus classified as severe acute respiratory syndrome coronavirus-2 (SARS-CoV-2) ^2^. A hallmark of severe COVID-19 is hyperinflammation ^3^, although cytokine expression patterns in individuals are diverse, leading to controversy over classification of COVID-19 related inflammation as cytokine storm, macrophage activation syndrome, multisystem inflammatory syndrome, etc. Regardless, inflammatory cytokines appear to play a principal role in mediating COVID-19 symptoms, therefore therapies which target these responses are paramount to treating severe COVID-19. As such, a fuller understanding of the cellular and molecular mechanisms mediating hypercytokinemia during SARS-CoV-2 infection is necessary.

Mononuclear phagocytes such as monocytes and macrophages are key constituents of the innate immune system, and produce pro-inflammatory cytokines during viral infection ^4–7^. Monocyte and monocyte-derived macrophage infiltration into the lungs has been linked to severe COVID-19 in single cell RNA sequencing studies ^8–11^ and postmortem analyses ^12–15^ in human patients, as well as during experimental infections in animal models including mice ^16,17^, hamsters ^18^, and various non-human primates ^19–23^. Monocytes in individuals infected with SARS-CoV-2 display phenotypic changes associated with hyperinflammation, including reduced HLA-DR expression ^24–26^, increased CD16 expression ^25,27–29^, and increased cytokine production ^30–33^. Both monocytes ^34–36^ and monocyte-derived macrophages ^37,38^ also produce pro-inflammatory cytokines under direct infection with SARS-CoV-2, although infection at least in macrophages appears to be abortive ^37,38^.

The past decade has seen an explosion in scientific interest in the regulation of immune cell activation and function by metabolic reprogramming. Under pro-inflammatory conditions, immune cells – including myeloid cells – generally undergo a switch to aerobic glycolysis which provides ATP sufficient to support cellular functions which propagate pro-inflammatory and anti-pathogen host responses ^39^. Recently, Codo *et al*. demonstrated pro-inflammatory glycolytic reprogramming in monocytes infected with SARS-CoV-2 ^34^, and SARS-CoV-2 also appears to alter monocyte lipid metabolism to promote lipid droplet formation which is associated with pro-inflammatory cytokine production ^35^.

SARS-CoV-2 therefore appears to reprogram metabolism in monocytes, but the viral factors which mediate these responses are unclear. Research in the 2003 epidemic SARS-CoV-1 suggested that the viral spike protein could mediate pro-inflammatory activation in macrophages ^40,41^, and recent evidence suggests the spike protein of SARS-CoV-2 also activates inflammatory responses in macrophages and monocytes both *in vitro* and *in vivo* ^42,43^. Given this, we hypothesized that spike protein binding to monocytes mediates glycolytic reprogramming to promote pro-inflammatory responses of these cells to SARS-CoV-2. Our results herein support this hypothesis, and we additionally report outcomes from experiments aimed at evaluating the responsible cellular signaling mechanisms, as well as potential pharmaceutical strategies for inhibiting these responses.

## Results

### Spike protein subunit 1 reprograms metabolism and promotes inflammatory responses

Recently it was demonstrated that SARS-CoV-2 promotes metabolic reprogramming in monocytes during infection ^34,35^. Research in SARS-CoV suggested that the viral spike protein induces inflammatory responses in macrophages ^40,41^, and this has recently been replicated using spike protein from SARS-CoV-2 ^42,43^. Likewise, spike protein binding to C-type lectins has recently been shown to mediate pro-inflammatory processes in myeloid cells ^44,45^. Therefore, we hypothesized that the SARS-CoV-2 spike protein mediates a pro-inflammatory metabolic reprogramming in monocytes which could be a basis for hypercytokinemia. Stimulation of isolated human classical monocytes with recombinant spike protein subunit 1 (S1) from SARS-CoV-2 induced glycolytic activation (Figure 1A) and suppressed oxidative phosphorylation (OXPHOS, Figure 1C) in a dose-dependent manner. The effect of S1 dose was significant for both extracellular acidification rate (F_2,14_=72.44, p<0.0001, Figure 1B) and oxygen consumption rate (F_2,14_=5.785, p=0.0147, Figure 1D) as measured by quantification of area under the response curve.

**Figure 1.**
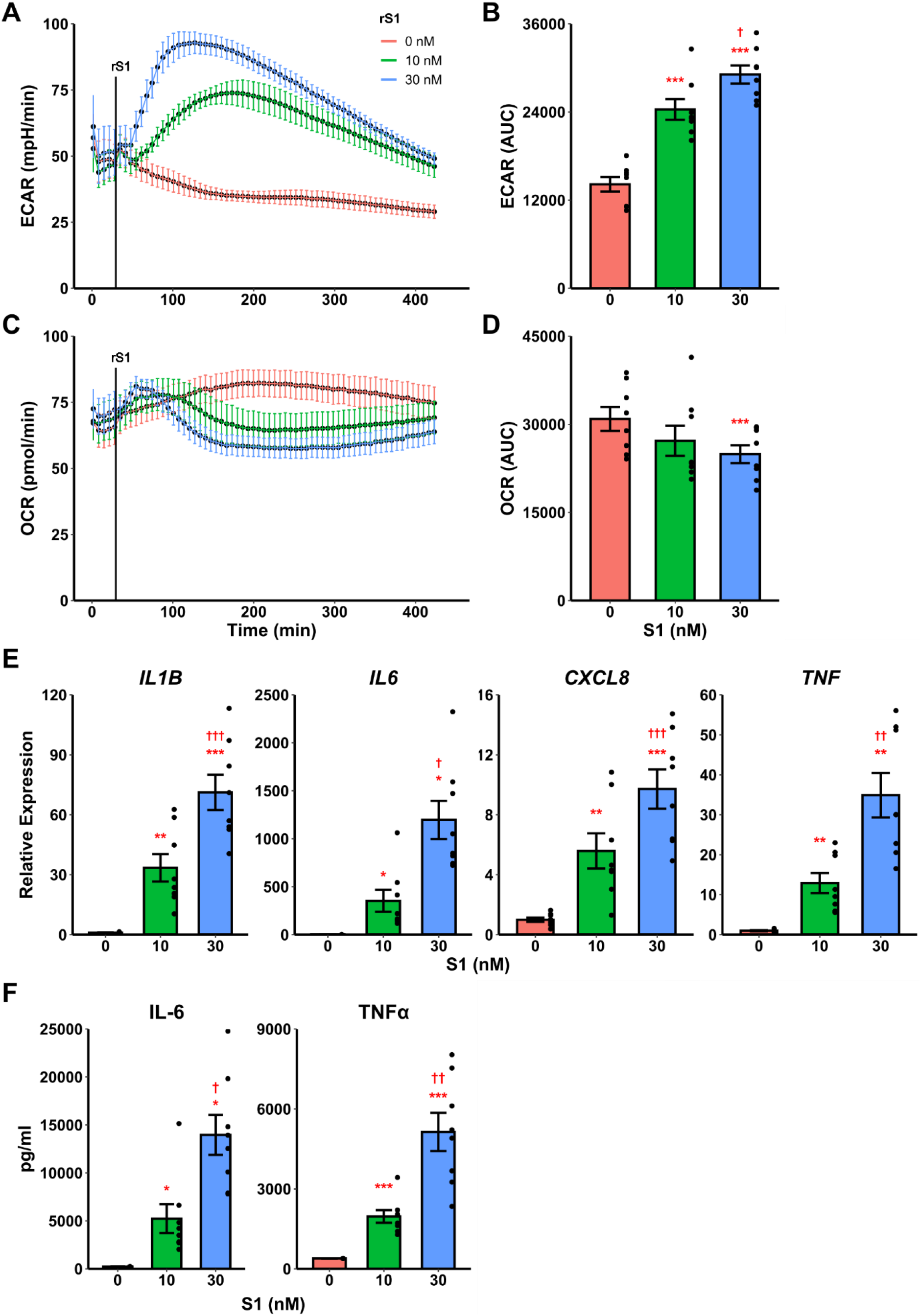
Recombinant SARS-CoV-2 spike protein subunit 1 (rS1) mediates immunometabolic activation of monocytes. (**A**) Monocytes increase extracellular acidification response rate (ECAR) in a dose-dependent manner when treated with rS1. (**B**) Quantification of ECAR by area under the curve (AUC). (**C**) rS1 treatment suppresses oxygen consumption rate (OCR) in monocytes in a dose-dependent fashion. (**D**) Quantification of OCR by AUC. (**E**) Gene expression analysis by qPCR reveals dose-dependent increases in responses of *IL1B, IL6, CXCL8*, and *TNF* to rS1 stimulation. (**F**) Protein expression analysis by ELISA reveals dose-dependent increases in responses of IL-6 and TNFα to rS1 stimulation. ECAR and OCR data in panels A-D are adjusted for values indexed to 1×10^5^ cells/well. *, **, ***: p<0.05, p<0.01, p<0.001 vs. 0 nM rS1. †, ††, †††: p<0.05, p<0.01, p<0.001 vs. 10 nM rS1. N=8 biological replicates.

Additionally, recombinant S1 treatment caused a dose-dependent increase transcription of pro-inflammatory cytokines (Figure 1E) including *IL1B* (F_2,14_=50.98, p<0.001), *IL6* (Friedman χ^2^_(df=2)_=16, p<0.001), *CXCL8* (F_2,14_=38.19, p<0.001), and *TNF* (F_2,14_=28.41, p<0.001) as measured by qPCR. These cytokines have been implicated in the pathogenesis of SARS-CoV-2 and in COVID-19-related hypercytokinemia in several studies ^25,46–51^. To confirm that increased transcription resulted in increased protein expression, we evaluated protein concentrations of key cytokines in the supernatant of S1-stimulated monocytes by enzyme-linked immunosorbent assay (ELISA) (Figure 1F). S1 increased protein expression of interleukin (IL)-6 (Friedman χ^2^_(df=2)_=16, p<0.001) and tumor necrosis factor (TNF)-α (F_2,14_=37.73, p<0.001) in a dose-dependent manner.

### Glycolytic response to spike protein is dependent on HIF-1α

Hypoxia inducible factor (HIF)-1α was demonstrated nearly 20 years ago to mediate pro-inflammatory responses in myeloid cells ^52^, and has more recently been shown to regulate glycolytic activation in monocytes, macrophages, and other immune cells ^53–55^. SARS-CoV-2 activates HIF-1α-mediated glycolysis in monocytes ^34^, so we reasoned that this was a likely downstream mechanism by which the viral spike protein causes this similar glycolytic reprogramming in our experiments. As above, treatment of monocytes with S1 activated glycolysis, and this effect was abrogated by pre-treatment with chetomin (Figure 2A/B, F_2,12_=42.43, p<0.001), which disrupts the interaction between HIF-1α and p300 to block the effects of the former ^56^. Pre-treatment with chetomin also strongly suppressed the cytokine response due to S1 treatment (Figure 2C), including blunting transcription of *IL1B* (F_2,12_=27.35, p<0.001), *IL6* (F_2,12_=16.11, p<0.001), *CXCL8* (F_2,12_=25.54, p<0.001), and *TNF* (F_2,12_=29.04, p<0.001). As such, HIF-1α appears to be a master regulator of both glycolytic reprogramming and inflammatory activation of monocytes under S1 stimulation.

**Figure 2.**
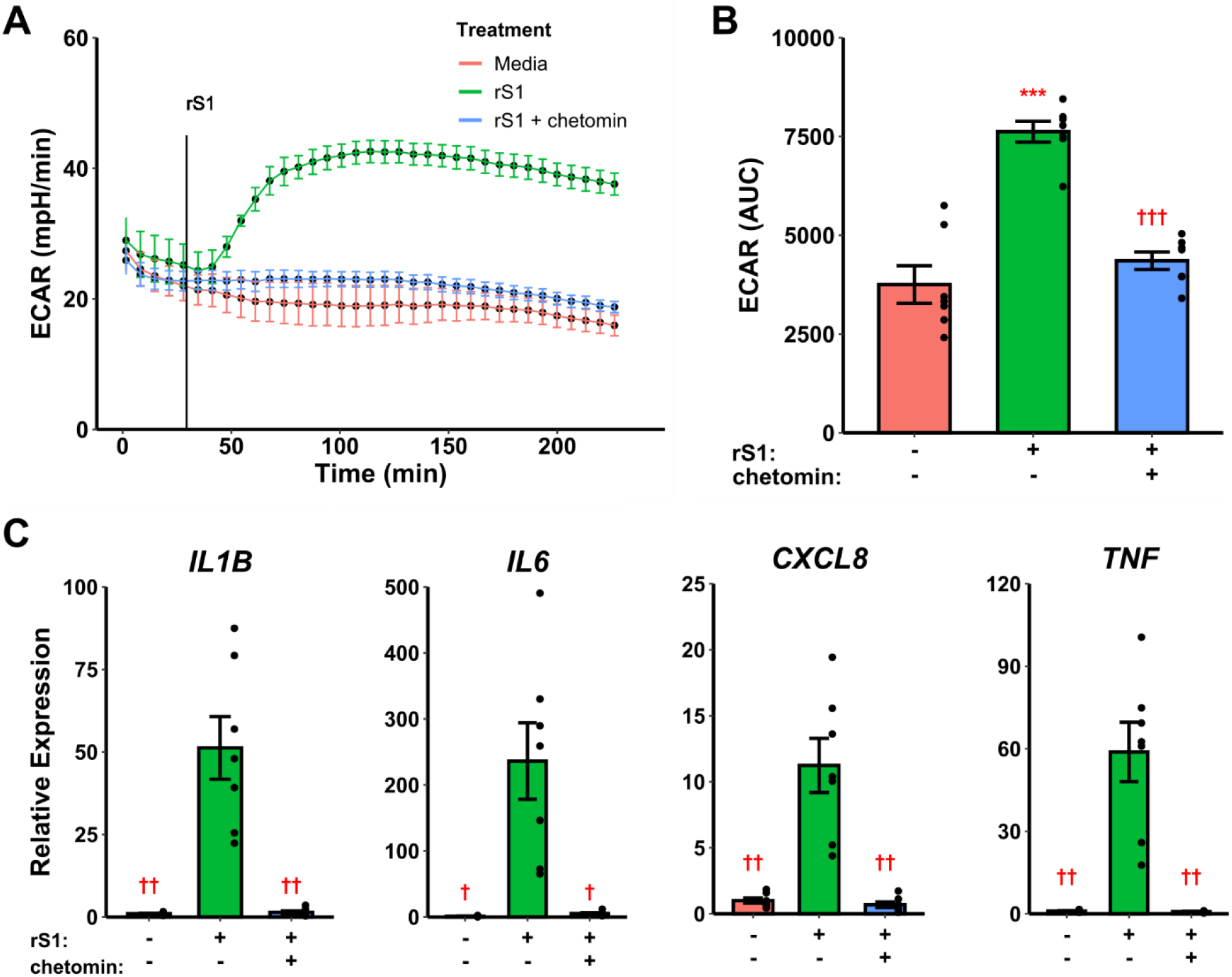
HIF-1α inhibition suppresses immunometabolic activation of monocytes due to recombinant spike protein (rS1). (**A**) Monocytes increase extracellular acidification response rate (ECAR) when treated with 30 nM rS1, but this is blocked by pre-treatment with chetomin. (**B**) Quantification of ECAR by area under the curve (AUC). (**C**) rS1 increase of expression of *IL1B, IL6, CXCL8*, and *TNF* is reversed by chetomin pre-treatment. *, **, ***: p<0.05, p<0.01, p<0.001 vs. untreated cells. †, ††, †††: p<0.05, p<0.01, p<0.001 vs. rS1-treated cells. N=7 biological replicates.

### Suppression of glycolysis alters inflammatory responses to spike protein

To determine whether metabolic reprogramming is responsible for altered cytokine responses to S1, we suppressed glycolytic responses during S1 treatment using 2-deoxyglucose (2-DG) pretreatment. Treatment of monocytes with 2-DG ablated monocyte glycolytic responses to S1 stimulation (Figure 3A) which was significant by comparison of area under the response curve (t_6_=−10.867, p<0.0001, Figure 3B). However, 2-DG also suppressed mitochondrial function in these cells (Figure 2C), though this was non-significant by area under the oxygen consumption (t_6_=−2.2284, p=0.0674, Figure 3D). This effect has been noted previously during responses to LPS ^57^. Anticipating this, we also included a condition where monocytes were cultured under glucose deprivation, as a second method of suppressing glycolytic activation. We noted a similar ablation of glycolytic responses to S1 using this strategy (Figure 3A) which was significant by area under the curve analysis (t_6_=−14.045, p<0.0001, Figure 3B). However, glucose deprivation caused an increase in oxygen consumption after S1 treatment (Figure 3C) which was significant compared to media- (t_6_=4.6618, p=0.0069) or 2-DG (t_6_=−15.607, p<0.001) pretreated monocytes (Figure 3D).

**Figure 3.**
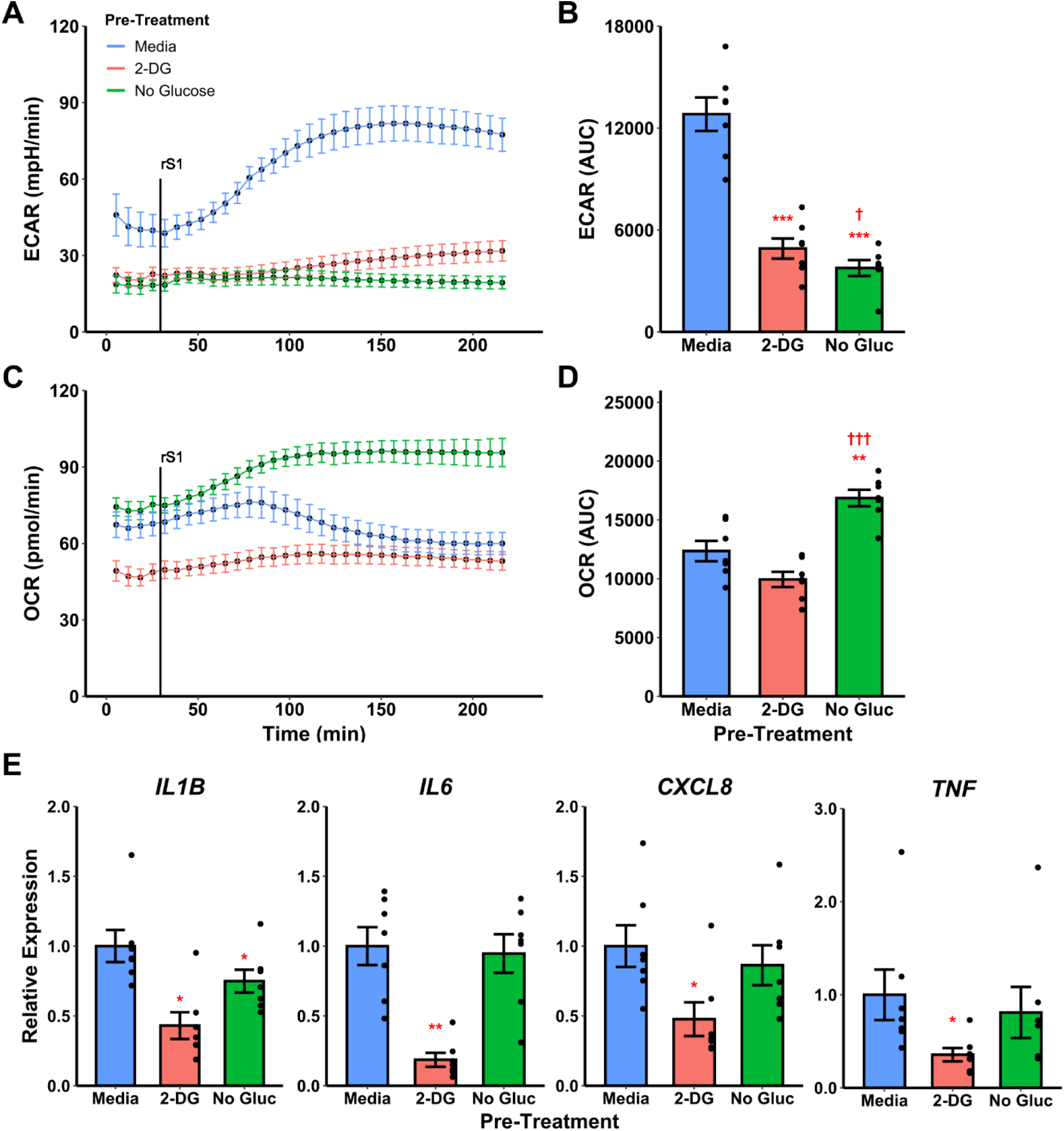
Targeting glycolysis has variable effects on recombinant spike protein (rS1) responses in monocytes. (**A**) Glucose deprivation or pre-treatment with 2-deoxyglucose (2-DG) block extracellular acidification rate (ECAR) increase due to rS1 treatment. (**B**) Quantification of ECAR by area under the curve (AUC). (**C**) 2-DG inhibits oxygen consumption rate in rS1-treated monocytes, but glucose-deprived monocytes upregulate OCR in response to rS1. (**D**) Quantification of OCR by AUC. (**E**) 2-DG blocks expression of *IL1B, IL6, CXCL8*, and *TNF* due to rS1 stimulation, but glucose deprivation has limited effects on cytokine expression. *, **, ***: p<0.05, p<0.01, p<0.001 vs. rS1-treated cells. †, ††, †††: p<0.05, p<0.01, p<0.001 vs. 2-DG-treated cells. N=7 biological replicates.

Pre-treatment of monocytes with 2-DG also strongly inhibited cytokine expression compared to cells treated with S1 (Figure 3E), including transcription of *IL1B* (*W*=0, p=0.0313), *IL6* (t_6_=−5.912, p=0.0021), *CXCL8* (*W*=0, p=0.0313), and *TNF* (*W*=0, p=0.0313). However, glucose deprived monocytes generally maintained their ability to transcribe pro-inflammatory cytokines in response to S1, with only *IL1B* expression showing a modest 25.1% reduction in glucose deprived compared to S1-treated monocytes (Figure 3E, *W*=0, p=0.0313). Monocytes appear to utilize fatty acid oxidation to compensate for loss of glycolysis during cytokine responses as has been previously demonstrated with LPS ^57–60^, and therefore the 2-DG-mediated suppression of S1-induced inflammation is likely due to its ability to suppress both glycolysis and mitochondrial metabolism in concert.

### AMPK controls the OXPHOS response to spike protein in the absence of glucose

Activation of OXPHOS appeared to compensate for loss of glycolysis under glucose deprivation conditions in S1-treated monocytes. Therefore, we examined the potential role of AMP-activated protein kinase (AMPK) in mediating this response. Compound C (dorsomorphin) is a selective inhibitor of AMPK ^61^, therefore we examined whether compound C would block OXPHOS activation by S1 in glucose deprived monocytes. Pre-treatment with compound C slightly reduced ECAR in glucose deprived monocytes compared to media-treated glucose deprived monocytes (Figure 4A/B), although this small effect is likely not biologically important. Compound C pre-treatment inhibited the OXPHOS response to S1 during glucose deprivation, but compound C-treated monocytes did not demonstrate suppressed OXPHOS as compared to S1-stimulated monocytes treated in the presence of glucose (Figure 4C/D, F_2,12_=25.81, p<0.001). Interestingly, compound C pre-treatment had a variable effect on cytokine gene expression, with only *TNF* expression significantly suppressed in glucose deprived monocytes treated with compound C compared to media (Figure 4E, Friedman χ^2^_(df=2)_=10.571, p=0.005). Expression of *IL1B, IL6*, and *CXCL8* were not significantly altered by compound C treatment, although *CXCL8* showed a trend toward increased expression in glucose deprived monocytes treated with compound C compared to media (p=0.0603).

**Figure 4.**
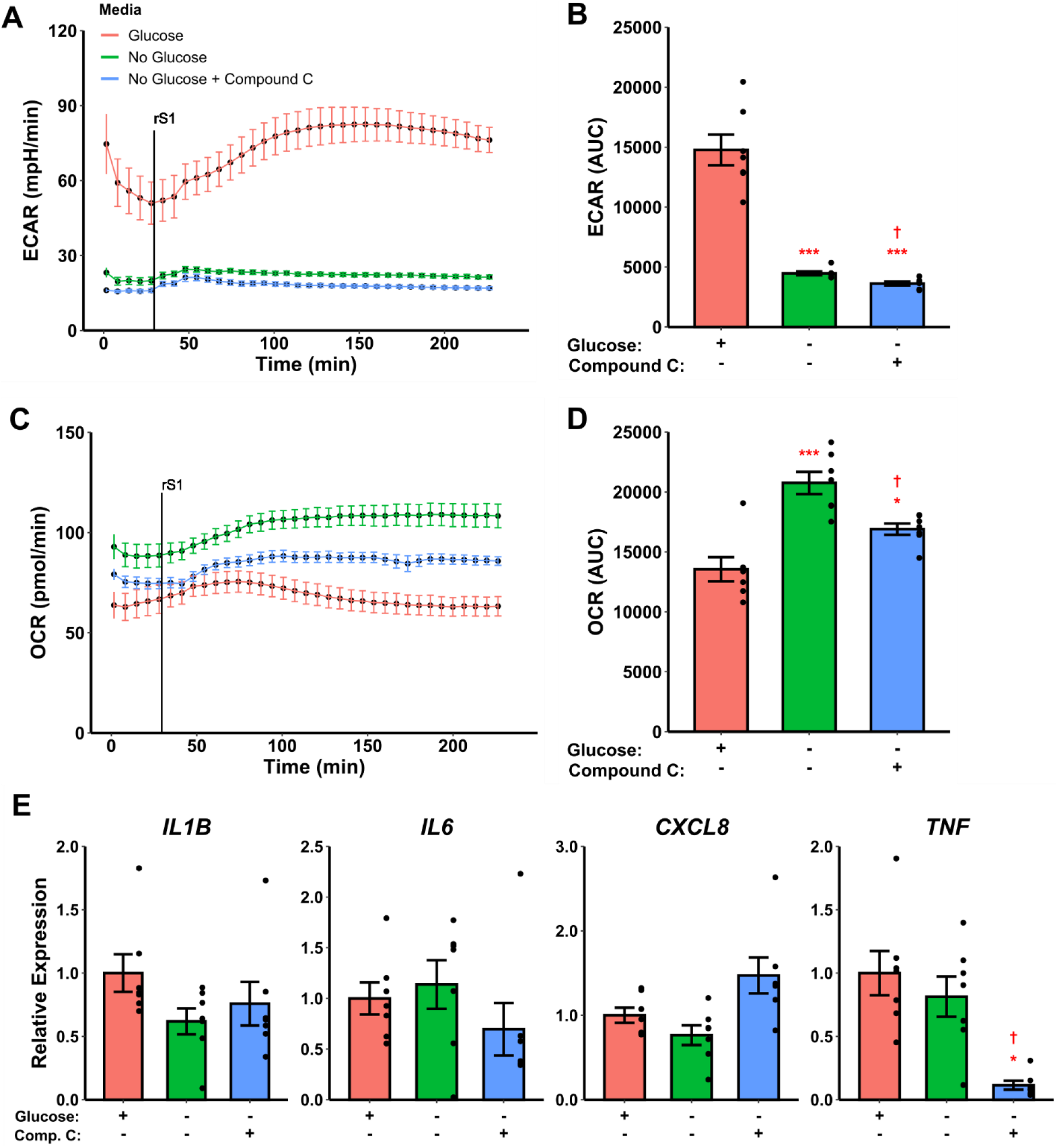
Inhibiting AMPK blocks metabolic response to recombinant spike protein subunit 1 (rS1) in glucose-deprived monocytes but has minimal effect on most cytokines. (**A**) Compound C pre-treatment has no effect on extracellular acidification (ECAR). (**B**) Quantification of ECAR by area under the curve (AUC). (**C**) Compound C blocks the increase in oxygen consumption rate (OCR) in rS1-treated glucose-deprived monocytes. (**D**) Quantification of OCR by AUC. (**E**) Compound C pre-treatment inhibits *TNF* expression in glucose-deprived monocytes stimulated with rS1, but does not significantly alter expression of *IL1B, IL6*, or *CXCL8*. *, **, ***: p<0.05, p<0.01, p<0.001 vs. glucose-treated cells. †, ††, †††: p<0.05, p<0.01, p<0.001 vs. glucose-deprived cells. N=7 biological replicates.

### Metformin abrogates inflammatory response to spike protein

The small molecule compounds chetomin and 2-deoxyglucose inhibited immunometabolic activation in monocytes, suggesting a potential strategy for treating hypercytokinemia during COVID-19. However, chetomin is not approved for use in humans, although it has shown efficacy *in vivo* in animal models ^56^. Additionally, 2-DG has poor efficacy in humans due to rapid metabolism and limited bioavailability ^62^. Therefore, we investigated the ability of the common diabetes and geroprotector drug metformin to inhibit cytokine production in S1-stimulated monocytes. Metformin activates AMPK ^61^ and (independently of AMPK) opposes the action of HIF-1α ^63,64^, and additionally inhibits mitochondrial metabolism through blocking complex I of the electron transport chain ^65,66^, thus we hypothesized that it would have a qualitatively similar effect to 2-DG in inhibiting cytokine production through dual inhibition of glycolysis and OXPHOS.

Pre-treatment with metformin abrogated the glycolytic response to S1 in monocytes (Figure 5A/B, F_2,12_=60.05, p<0.001) and strongly inhibited cellular respiration (Figure 5C/D, Friedman χ^2^_(df=2)_=12.286, p=0.0021) in Seahorse assays. Likewise, metformin pre-treatment suppressed cytokine responses to S1 treatment in monocytes (Figure 5E), including *IL1B* (Friedman χ^2^_(df=2)_=12.286, p=0.0021), *IL6* (Friedman χ^2^_(df=2)_=10.571, p=0.0051), *CXCL8* (F_2,12_=68.18, p<0.0001), and *TNF* (Friedman χ^2^_(df=2)_=12.286, p=0.0021).

**Figure 5.**
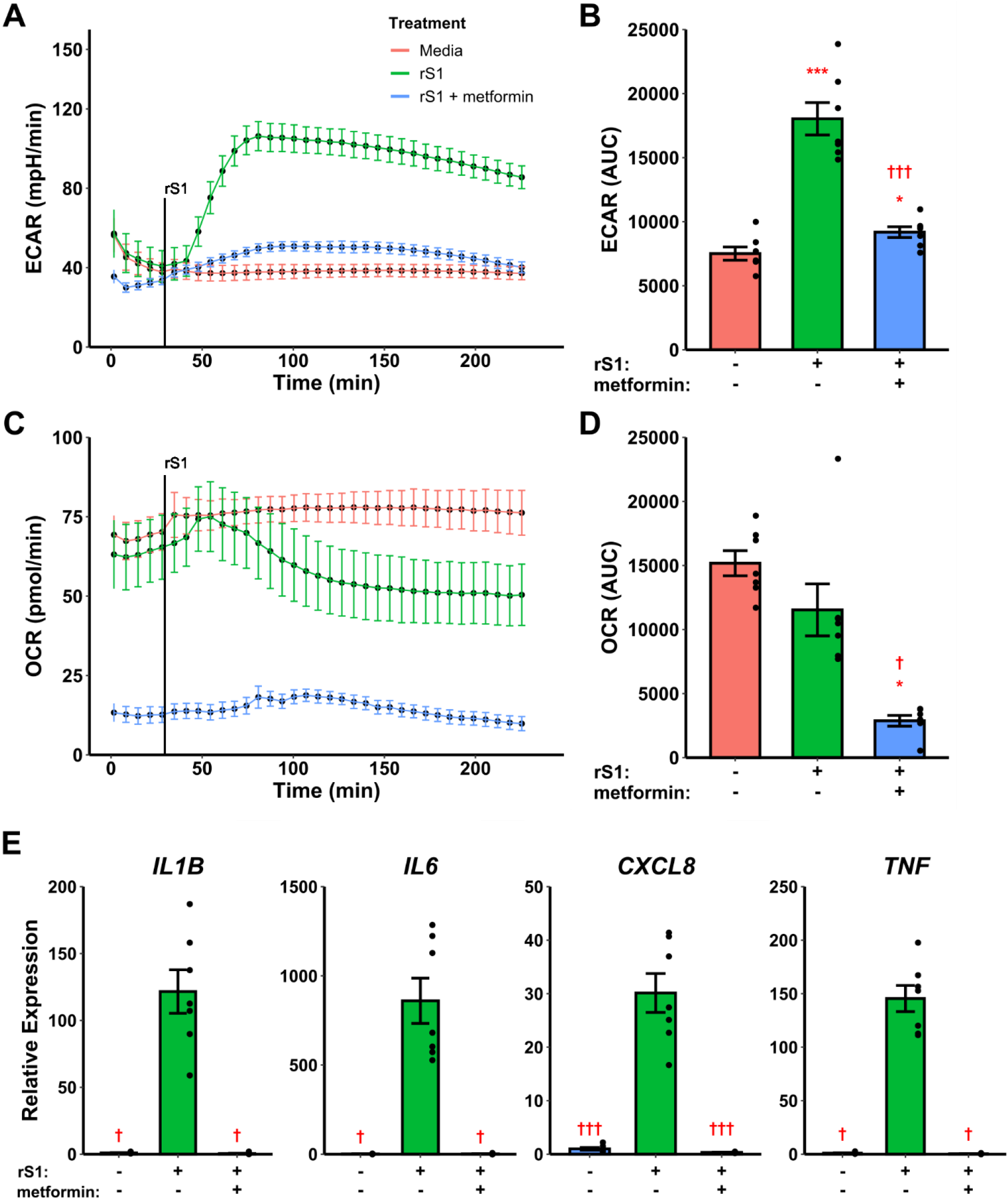
Metformin suppresses immunometabolic activation in monocytes treated with recombinant spike protein (rS1). (**A**) Metformin pre-treatment blocks the increase in extracellular acidification rate (ECAR) mediated by rS1. (**B**) Quantification of ECAR by area under the curve (AUC). (**C**) Metformin suppresses oxygen consumption rate (OCR). (**D**) Quantification of OCR by AUC. (**E**) Metformin suppresses cytokine responses, as demonstrated by gene expression of *IL1B, IL6, CXCL8*, and *TNF*, during rS1 stimulation in monocytes. *, **, ***: p<0.05, p<0.01, p<0.001 vs. unstimulated cells. †, ††, †††: p<0.05, p<0.01, p<0.001 vs. rS1-treated cells. N=7 biological replicates.

### Metformin abrogates IL-6 production in virus-stimulated monocytes

Recent evidence suggests that myeloid cells recognize SARS-CoV-2 spike protein through C-type lectins ^44,45^. However, the SARS-CoV-2 virion also contains additional immunoregulatory and pro-inflammatory proteins ^67,68^, therefore we examined the ability of metformin to block cytokine responses to live SARS-CoV-2. Monocytes treated for 24 hr with SARS-CoV-2 increased expression of IL-6 protein, and this was suppressed by metformin pre-treatment (Figure 6, F_2,14_=11.48, p=0.0011), suggesting that the anti-inflammatory effect of metformin is not specific to spike protein-stimulated cytokine responses in monocytes.

**Figure 6.**
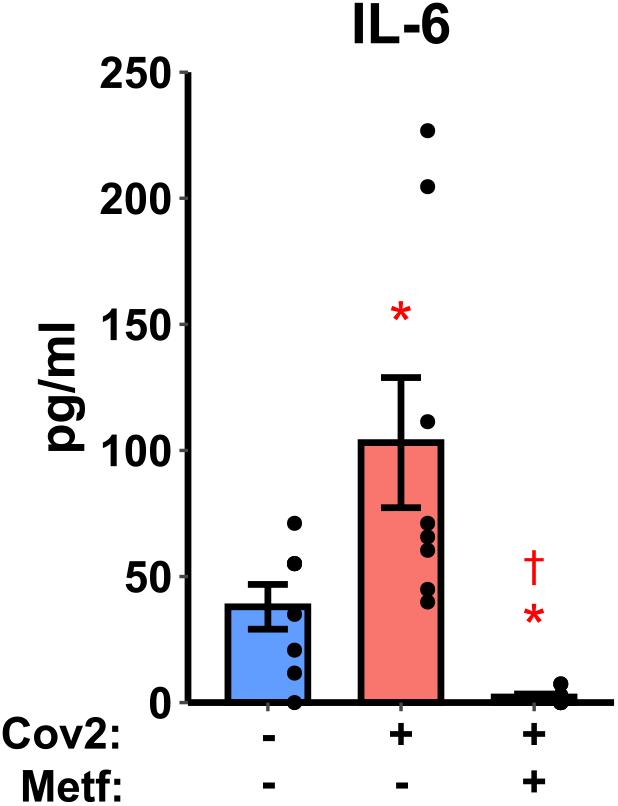
Metformin inhibits IL-6 production in monocytes infected with SARS-CoV-2 strain WA1/2020 (Cov2) at 0.5 MOI. *: p<0.05 vs. uninfected cells. †: p<0.05 vs. Cov2-infected cells. N=8 biological replicates.

## Discussion

The present study resulted in several advances of major importance for the understanding of SARS-CoV-2 innate immune responses. First, we report here that monocytes treated with recombinant spike protein subunit 1 from the current pandemic SARS-CoV-2 undergo a dose-dependent increase in glycolysis which is controlled by HIF-1α and mediates the production of pro-inflammatory cytokines. These data suggest an initial signaling event which precipitates changes in glucose and lipid metabolism during SARS-CoV-2 infection in monocytes which have been previously reported to be linked to inflammatory activation ^34,35^. Monocyte and monocyte-derived macrophages are substantially enriched in the lungs of SARS-CoV-2-infected individuals with severe COVID-19 ^8,9,12–15^ and respond to experimental viral infection by producing pro-inflammatory cytokines ^34–38^, therefore these results reflect a potential mechanism by which hypercytokinemia occurs during the early innate immune response to SARS-CoV-2.

Importantly, the available evidence suggests that infection of monocytes/macrophages by SARS-CoV-2 is abortive ^37,38,69^, thus recognition of SARS-CoV-2 structural proteins or genomic material is the likely mechanism by which direct infection precipitates inflammatory responses in this cell type. Our data suggest the spike protein is one such determinant, although we cannot conclude it is the only such mechanism given that recent reports have demonstrated inflammatory responses in macrophages treated with the SARS-CoV-2 envelope protein ^67^. It is also possible, however, that direct viral binding to monocytes is not the only way in which these cells can be exposed to the viral spike protein. Recent evidence suggests that vaccine antigens including S1 are released into the circulation following vaccination ^70^, and this represents a potential pro-inflammatory stimulus for monocytes. Monocyte/macrophage recognition of S1 may also contribute to the local (muscle) inflammatory response during vaccination. Additionally, the viral spike protein undergoes cleavage by furin during binding to ACE2 ^71^, and it has been suggested that this could lead to release of the S1 subunit ^72^, although to date this is speculative.

ACE2 has limited expression on immune cells including monocytes and macrophages ^73^, which has called into question whether they can directly recognize SARS-CoV-2. In this study we did not identify the mechanism for monocyte recognition of S1, but several recent papers have shed light on this. Two reports recently demonstrated spike binding to C-type lectin receptors ^44,45^ which mediates pro-inflammatory signaling in myeloid cells. Likewise, monocytes and macrophages express high levels of CD147 ^73^, and this receptor has been shown to recognize spike protein and contribute to activation of T cells ^74^. Monocytes therefore have multiple methods of recognizing S1, and the receptor(s) responsible for signaling to induce immunometabolic activation deserve further investigation.

The second major advance in this study is the identification of metformin as a potential immunometabolic regulator of inflammatory responses to SARS-CoV-2. Small molecule inhibitors of HIF-1α (chetomin) and glucose metabolism (2-deoxyglucose) blocked cytokine production in S1-treated monocytes, suggesting that interfering with downstream signaling pathways activated by spike protein binding is a potential therapeutic strategy to target inflammation during COVID-19. As these compounds are not approved for human use or have low efficacy in humans as described above, we evaluated the ability of metformin to suppress glycolytic reprogramming and cytokine production in S1-stimulated monocytes. Metformin reduced cytokine production and strongly inhibited both glycolysis and cellular respiration in culture, suggesting it as a potential treatment for hyperinflammation during COVID-19. Further, metformin blocked IL-6 production in monocytes infected with live SARS-CoV-2, suggesting this effect is not limited to artificial stimulation conditions with purified recombinant protein.

Metformin is extremely inexpensive compared to many pharmaceuticals, with an estimated manufacturing cost under 10 USD per kg for the active ingredient ^75^ and a monthly wholesale cost as low as 25 USD ^76^. Metformin has been previously noted as a treatment for non-COVID acute respiratory distress syndrome ^77^ and is a potent suppressor of immune activation of monocytes and macrophages by other molecules including LPS ^78–80^. Additionally, several epidemiological studies have noted decreased mortality ^81–85^ and inflammation ^86,87^ in COVID-19 patients who were taking metformin prior to diagnosis. Therefore, given these observations and its low cost, excellent safety profile, wide availability, and efficacy in inhibiting inflammatory responses to S1 *in vitro*, metformin is a promising candidate for further exploration as a COVID-19 therapeutic. Our study is limited to a single *in vitro* measure of metformin as a therapeutic for COVID-19, so a great deal of further study is necessary in order to establish this drug as a viable treatment.

### Conclusion

In summary, we demonstrate here that the spike protein subunit 1 from SARS-CoV-2 causes activation of HIF-1α dependent glycolysis and inflammatory cytokine production in monocytes which can be suppressed by treatment with the diabetes drug metformin. These experiments detail a mechanism by which SARS-CoV-2 mediates metabolic reprogramming previously described in human monocytes, and additionally provides a potential mechanism for the observation that metformin is protective against mortality in COVID-19 patients. Continued research in this area has the potential to define therapeutic strategies and additional molecular targets for the treatment of COVID-19-associated hyperinflammation.

## Methods

### Subjects

Healthy 18–35-year-old subjects (N=14) were recruited without respect to sex or race. Participants reported to the laboratory approximately every two weeks for blood collection, and 8-24 ml blood was collected into EDTA-treated vacutainer tubes by venipuncture. Blood was immediately used for cell isolations as described below. All human subjects activities were approved by the Institutional Review Board at the University of Memphis under protocol 4316, and subjects provided informed consent prior to enrollment.

### Cell isolations

Assays were performed on purified human classical monocytes isolated using immunomagnetic negative sorting (EasySep Direct Human Monocyte Isolation Kit, StemCell Technologies, Cambridge, MA). As we have previously described ^88^, this procedure results in a highly pure (> 85%) population of classical monocytes, with depletion of intermediate and non-classical monocytes due to the presence of an anti-CD16 antibody in the cocktail. Isolation purity was verified at several points throughout the current study and averaged approximately 90% (not shown). Cells were counted at 10× dilution using a Scepter cell counter (Millipore Sigma, St. Louis, MO). Isolated monocytes were immediately utilized in downstream assays, and no cells were frozen for later use.

### Media and reagents

Unless otherwise specified, all assays were performed using Seahorse XF base DMEM medium (Agilent, Santa Clara, CA) supplemented with 10 mM glucose and 2 mM L-glutamine (Millipore Sigma, St. Louis, MO). Assays utilizing glucose deprivation omitted glucose from the media preparation. Media was not supplemented with fetal bovine serum or other additives. Recombinant spike protein subunit 1 (S1) was purchased from RayBiotech (Peachtree Corners, GA). 2-deoxyglucose, chetomin, compound C, and metformin were purchased from Millipore Sigma (St. Louis, MO). SARS-CoV-2 WA1/2020 strain was provided by Dr. Colleen Jonsson, Regional Biocontainment Laboratory, University of Tennessee Health Science Center.

### Seahorse extracellular flux

Glycolysis and oxidative phosphorylation were respectively quantified via kinetic monitoring of extracellular acidification rate (ECAR) and oxygen consumption rate (OCR) on a Seahorse XFp analyzer (Agilent, Santa Clara, CA). For all assays, monocytes were plated at 1.5×10^5^ cells per well, and wells A and H of the XFp plate were background wells with no cells. All analyses were run in duplicate. Plated cells were incubated at 37°C in a non-CO_2_ incubator for 1 hour prior to assays to stabilize pH. All wells were imaged at 10× magnification for cell counting in order to adjust raw measurements for cell number.

For quantification of dose response to S1, 5 basal measurements were made, followed by injection of media (wells B-C), 100 nM spike protein (wells D-E), or 300 nM spike protein (wells F-G). After injection into existing media in the well, spike protein concentrations were 10-fold lower than injection concentrations, thereby giving final spike protein concentrations of 0 nM, 10 nM, or 30 nM. Following injection, ECAR and OCR were monitored serially for 60 measurements. Following the assay, cell culture supernatants were removed, pooled by duplicate, and stored at −80°C. Cells were then lysed with 100 μl Trizol (Thermo Fisher Scientific, Waltham, MA), pooled by duplicate, and stored at −80°C as we have previously described ^89^.

For chetomin and metformin Seahorse assays, cells were incubated in media as above (wells B-E), or either 10 nM chetomin or 50 mM metformin during the 1-hour pre-incubation period (wells F-G). 5 basal ECAR/OCR measurements were performed, followed by injection of media (wells B-C) or 300 nM spike protein (wells D-G) for a final concentration of 0 nM (wells B-C) or 30 nM spike protein (wells D-G) as above. Following injection, ECAR and OCR were monitored serially for 30 measurements. Cell culture supernatants and Trizol lysates were processed as described above following the end of the assay.

For glycolysis inhibition assays, cells were incubated in media, 10 mM 2-deoxyglucose, or media without glucose (glucose deprivation) during the 1-hour pre-incubation period. 5 basal ECAR/OCR measurements were performed, followed by injection of 300 nM spike protein to all wells for a final concentration of 30 nM spike protein per well as above. Spike protein was prepared in non-glucose media for the glucose deprivation condition. Following injection, ECAR and OCR were monitored serially for 30 measurements. Cell culture supernatants and Trizol lysates were processed as described above following the end of the assay.

For glycolysis inhibition with compound C assays, cells were incubated in media, glucose deprivation media, or glucose deprivation media plus 10 μM compound C during the 1-hour pre-incubation period. 5 basal ECAR/OCR measurements were performed, followed by injection of 300 nM spike protein to all wells for a final concentration of 30 nM spike protein per well as above. Spike protein was prepared in non-glucose media for the glucose deprivation condition. Following injection, ECAR and OCR were monitored serially for 30 measurements. Cell culture supernatants and Trizol lysates were processed as described above following the end of the assay.

### SARS-CoV-2 infections

Isolated monocytes were incubated in RPMI-1640 media (Gibco, Thermo Fisher Scientific, Waltham, MA) supplemented with 10% fetal bovine serum (Gibco), with or without 50 mM metformin, for 1-hour. Cells were then treated with media or infected with SARS-CoV-2 virus (WA1/2020 isolate) at 0.5 MOI and incubated for 24 hours. Cell culture supernatants were collected from untreated and infected cells and stored at −80°C until analysis.

### Gene and protein expression analysis

RNA isolation was performed using the Trizol procedure based on manufacturer’s instructions from cells lysed directly in the microplate or Seahorse plate wells as applicable. Isolated RNA (300-400 ng depending on experiment) was reverse-transcribed to cDNA using a High-Capacity cDNA Reverse Transcription Kit (Thermo Fisher Scientific, Waltham, MA). Gene expression was analyzed using commercial pre-validated gene expression assays and Taqman reagents (Thermo Fisher Scientific, Waltham, MA). Relative gene expression was quantified using the 2^-ΔΔCt^ method ^90^ against *B2M* or *ACTB* as housekeeping genes. Primer/probe IDs were: *B2M* Hs00187842_m1; *ACTB* Hs03023943_g1; *IL1B* Hs01555410_m1; *IL6* Hs00174131_m1; *CXCL8* Hs00174103_m1; *TNF* Hs00174128_m1.

For protein quantification, cell culture supernatants harvested from microplates or Seahorse XFp plates were analyzed via ELISA. Commercial DuoSet matched-antibody reagent sets were purchased from R&D Systems (Minneapolis, MN) for quantifying human IL-6 and human TNFα and used according to manufacturer’s instructions. All samples were run in duplicate at 5× dilution (SARS-CoV-2 assays) or 50× dilution (Seahorse S1 dose response assays) and assessed against a standard curve.

Protein concentration of angiotensin converting enzyme 2 (ACE2) and C-reactive protein (CRP) was performed by ELISA on plasma samples collected by venipuncture from subjects at the beginning of the study. Peripheral blood was collected by venipuncture into EDTA-coated vacutainer tubes, centrifuged at 1,500×*g* for 15 min, aliquoted, and stored at −80°C until analysis. Plasma samples were analyzed in duplicate at 10× (ACE2) or 10,000× (CRP) using commercial DuoSet matched-antibody reagent kits (R&D Systems) according to manufacturer’s instructions and assessed against a standard curve.

### Data Processing and Statistical Analysis

All data processing and statistical analyses were performed using R v. 3.6.2 ^91^. Isolated monocytes from each subject were given all treatments for each experiment, so data were paired and analyzed using within-subjects designs. Data were checked for normality by Shapiro-Wilk test and analyzed by one-way repeated measures ANOVA (RM-ANOVA, for data which met the normality assumption) or Friedman’s test (for data which did not meet the normality assumption). For analyses with significant main effects, post hoc mean separation was performed using pairwise paired T tests (for RM-ANOVA) or pairwise Wilcoxon signed-rank tests (for Friedman’s tests) with p-value adjustment using the Holm-Bonferroni method ^92^. Significance cutoff was p<0.05.

## Acknowledgements

The authors would like to acknowledge the participants in this study. The authors thank Jyothi Parvathareddy and Colleen Jonsson from the Regional Biocontainment Laboratory at the University of Tennessee Health Science Center for providing SARS-CoV-2 virus stocks.

## Authors’ Contributions

BDP conceived the study. TJC and BDP designed experiments. TJC, RSE, JRY, KLD, and BDP collected data.

BDP analyzed data and prepared the first manuscript draft. TJC, RSE, JRY, KLD, and BDP edited the manuscript draft. All authors read and approved the final manuscript.

